# Bioinformatic analysis of human ZPR1 gene pathogenic exome mutations

**DOI:** 10.1101/2024.02.27.582196

**Authors:** Jeremiah I. Abok, William S. Garver, Jeremy S. Edwards

## Abstract

Advanced sequencing technologies enable rapid detection of sequence variants, aiming to uncover the molecular foundations of human genetic disorders. The challenge lies in interpreting the influence of new exome variants that lead to diverse phenotypes. Our study introduces a detailed, multi-tiered method for assessing the impact of novel variants, particularly focusing on the zinc finger protein 1 (ZPR1) gene. Herein, we employed a combination of variant effect predictors, protein stability analyses, and the American College of Medical Genetics and Association of Molecular Pathology (ACMG/AMP) guidelines. Our structural analysis pinpoints specific amino acid residues in the ZPR1 zinc finger domains that are sensitive to changes, distinguishing between benign and disease-causing coding variants using rigorous *in silico* tools. We examined 223 germline ZPR1 exome variants, uncovering significant ethnic disparities in the frequency of heterozygous harmful ZPR1 variants, ranging from 0.04% in the Ashkenazi Jewish population to 0.34% in African/African Americans. Additionally, the discovery of three homozygous carriers in European and South Asian groups suggests a higher occurrence of ZPR1 variants in these demographics, meriting further exploration. This research provides insights into the prevalence and implications of amino acid substitutions in the ZPR1 protein.

## Introduction

The zinc finger protein 1 (ZPR1) gene in humans produces a protein comprising 459 amino acids. This protein is recognized as isoform I and represents the primary version of ZPR1 [1]. ZPR1 is a C4-type zinc finger protein and part of a family of zinc finger proteins linked to various conditions, including cancer, non-alcoholic fatty liver disease (NAFLD), type 2 diabetes mellitus (T2DM), and other genetic disorders [2]. The gene’s cytogenetic location is on chromosome 11 at the specific site 11q23.3. Its coordinates are Chr11: 11:116,773,799-116,788,023 on the reverse strand, according to the GRCh38.p13 [3].

In 2018, Ito et al. reported a novel rare autosomal recessive disorder (RARD) observed among four children in three New Mexican Hispanic ancestral families. A proband from one of these families had an exome sequence that showed a homozygous c.587T>C (p.Ile196Thr) mutation in the ZPR1 gene [4]. Further analysis revealed that this mutation was present in a heterozygous state among the parents and unaffected siblings [4]. From a clinical perspective, patients with homozygous mutations exhibited a systemic syndrome marked by growth limitations both before and after birth, inborn hair loss, kidney dysfunction, developmental delays, hearing impairment, and increased early mortality. In another family, it was found that the parents of an affected individual were heterozygous carriers of the same ZPR1 mutation, and there were no other siblings with a homozygous manifestation of the condition [4]. The fact that both families resided in the Rio Grande Valley area of New Mexico suggests the possibility of a founder effect mutation [4] combined with endogamous practices [5, 6].

It is thought that the function of the ZPR1 gene is similar to other genes linked to primordial dwarfism, which negatively impacts cell cycle progression and cell proliferation [4,7]. For example, mutations present in the ORC1, ORC4, ORC6, CDT1, and CDC6 genes alter G1/S transition and S phase progression by perturbing the pre-replication complex and predisposition to Meier-Gorlin syndrome [8, 9], and mutations present in the ATR gene alter S phase progression and predisposition to Seckel syndrome type 1 disease [10, 11]. The missense mutation in the ZPR1 gene c.587T>C (p.Ile196Thr) has been shown to stop the cell cycle from moving past the G1 phase [4], which is similar to genes linked to primordial dwarfism. At least two studies have indicated that ZPR1 protein primarily resides in the cytoplasm of quiescent mammalian cells but is translocated into the nucleus after binding eukaryotic translation elongation factor 1A (eEF1A), a process regulated by epidermal growth factor (EGF) and disrupts the interaction with epidermal growth factor receptor (EGFR) [12, 13]. Another study indicated that the ZPR1 protein binds to the cytoplasmic tyrosine kinase domain of EGFR via two zinc finger structural motifs and accumulates in the nucleolus of proliferating cells [14]. Collectively, these findings imply that the ZPR1 protein plays a crucial role in the nucleolar function of proliferating cells. [13].

It has been reported that decreased expression of ZPR1 in humans contributes to spinal muscular atrophy (SMA), a neurodegenerative condition. In cells deficient in ZPR1, the nuclear positioning of survival motor neurons (SMN) and the NPAT transcription factor is disturbed, leading to a blockage in S phase progression and arrests in both the G1 and G2 phases. [15]. These changes in subnuclear architecture and cell cycle progression could be attributed to transcriptional defects [15]. In normal proliferating cells, the ZPR1 protein spreads throughout the cell during the G1 and G2/M phases and redistributes to the nucleus during the S phase [16]. The eEF1A has a conserved binding epitope required for normal cell growth, proliferation, and cell cycle progression. Based on a previous study [17], the ZPR1 protein can form a complex with the GDP-bound eEF1A to regulate the activation of eEF1A. Also, it can form a complex with the SMN protein in response to stimuli induced by EGF. The ZPR1 domains are structurally different from each other, which leads to functional divergence that changes how the ZPR1 protein interacts with those two complexes [17]. Nonetheless, the truncated mutant ZPR1-ΔA (193–246) can still bind eEF1A in the absence of Zn2+ [18]. To date, the interactions between ZPR1 and eEF1A2 continue to represent an incredibly critical research area concerning motor neuron biology [19]. A study has indicated that the ZPR1 protein is capable of binding to RNA polymerase II to increase the expression of the SMN2 gene [20], and overexpressing the ZPR1 protein can raise the levels of SMN throughout the body and prevent the disease from starting in SMA mice [20].

Besides the previously mentioned RARD linked to ZPR1 deficiency, a prevalent mutation, rs964184, has related to abnormal glucose metabolism and Type 2 Diabetes Mellitus (T2DM) in Japanese populations [21]. Another frequent mutation, rs2075294, located in the ZPR1 gene within the APOA1/C3/A4/A5-ZPR1-BUD13 gene cluster, is associated with dyslipidemia [22]. Supporting these observations, a separate study [2] showed that the ZPR1 protein interacts with high-fat diets, influencing energy metabolism and neurological degeneration. Furthermore, ZPR1 expression levels in breast cancer tissues were significantly elevated compared to adjacent non-tumorous tissues (P < 0.001), suggesting a potential role of ZPR1 gene mutations in promoting the spread of breast cancer through cell invasion and migration, possibly via the ERK/GSK3/Snail signaling pathway [23]. Finally, a reduction in ZPR1 gene expression is linked with the development of non-small cell lung cancer (NSCLC), indicating that the ZPR1 protein might suppress proliferation and invasion of NSCLC cells by inhibiting the FAK-AKT signaling pathway [24].

Since the ZPR1 protein has a significant role in multiple human diseases, it is important to know the functional effects of the potential pathogenic mutations in the gene. We looked at the ZPR1 pathogenic exome mutations in a large-scale quantitative population genetics and ethno-geographic way in this study. Using the data from the Genome Aggregation Database (gnomAD), a large publicly accessible database of population variation derived from harmonized sequencing data, we predicted the effects of 223 rare ZPR1 exome mutations using four computational methods: SIFT, Polyphen-HVAR, FoldX, and Consurf. The obtained results have improved our understanding of the molecular basis of how the ZPR1 gene or protein, and the functional-related binding partner proteins interact and predispose to disease susceptibility.

## Materials and Methods

### Subjects of analysis and data availability

The exome data of the dataset GRCH37/hg19 used in this study is publicly and openly available in the Genome Aggregation Database V 2.1.1 at https://gnomad.broadinstitute.org/. The data was collected on March 12, 2021.

### Bioinformatic analyses of ZPR1 gene mutations

ANNOVAR [25], a software package consisting of a series of bioinformatic tools, was used in the bioinformatic analyses. Specifically, to annotate the functions of the ZPR1 gene mutations, the script annotate_variation.pl was used with GRCH37/hg19. The gnomAD allele frequency for exome variants was used for gnomad_exome. For the impact scores for the mutations, the script table_annovar.pl was used to run two variant effect predictor (VEP) algorithms: SIFT [26] and Polyphen2_HVAR [27]. To achieve a stringent classification of the pathogenic and neutral mutations, the default thresholds of the calculated scores were adjusted based on a previous study [26, 27]. For SIFT, a score of 0.50–1.00 and a score of 0.0–0.49 were set for neutral and pathogenic mutations, respectively. For polyphen2_HVAR, a score of 0.00–0.89 and a score of 0.91–1.00 were set for neural and pathogenic mutations, respectively. We used ten more VEP algorithms: SIFT4G[40], Polyphen2_HDIV[27], LRT[41], MutationTaster[42], MutationAssessor[43], PROVEAN[44], M-CAP[33], BayesDel [34], ClinPred [35], and FATHMM-MKL [36] to look into the effects of the pathogenic mutations found by the SIFT and Polyphen2_HVAR scores. Scores from each of these ten VEP algorithms were calculated using the script table_annovar.pl with the dbnsfp41a keyword specification. To be aware of, this ANNOVAR analysis did not include exome mutations that gnomAD had flagged. All the encoded protein products involved in the present study correspond to isoform I. Mutations leading to a different isoform were not included.

### Protein evolutionary conservation analysis

The ConSurf server (https://consurf.tau.ac.il/) [28] scores the evolutionary conservation based on an empirical Bayesian algorithm and the phylogenic relations between close homologous amino acid sequences that are calculated with the Maximum Likelihood (ML) method [29]. It was used to analyze the evolutionary conservation of the ZPR1 protein. In this analysis, a conservation score of 1–9 was obtained for each residue position. The positions were classified based on the scores as fluid (1–4), intermediary (5–6), or highly conserved (7–9). Based on recommendations from the previous studies, a residue at a position that is highly conserved and exposed was proposed to be functional; a residue at a position that is highly conserved but buried was proposed to be structural.

### Structural analysis of the ZPR1 protein

Iterative Threading Assembly Refinement (I-TASSER) [30] was used to model the structure of the ZPR1 protein with no specific assumptions. The modeling conducted with I-TASSER is based on a fold recognition that meta-threads the amino acid sequence of ZPR1 against the PDB (Protein Data Bank) library to identify the homolog template(s). The major threading template I-TASSER identified in this modeling is Mus musculus ZPR1 (PDBID: 2qkdA) [31]. The structural stability of the ZPR1 protein resulting from the gene mutations was predicted by calculating the free energy change (ΔΔG) of folding using the software FoldX 5.0[49].

### Statistical analysis

A Chi-square test using one-sided P-values was performed to determine the significant differences between variant frequencies and gender distributions. A resultant value less than 0.05 was considered to indicate a significant difference.

### Compliance with ethical standards

Written informed consent was obtained from each participant before participation, in accordance with the Declaration of Helsinki.

## Results

### Identification of the ZPR1 pathogenic exome mutations

The gnomAD dataset, comprising 141,456 unrelated individuals, was created through extensive exome and whole genome sequencing, aligning with the GRCH37/hg19 reference. This dataset encompasses exome data from seven major ethnic groups, including Europeans (67,709 individuals), Latino/Americans (17,296), South Asians (15,308), East Asians (9,197), Africans/African Americans (8,128), Ashkenazi Jews (5,040), and a miscellaneous group (3,070). The dataset, excluding the ‘other’ category, totals 122,678 individuals from six specific ethnic groups, providing a rich resource for population-genetic and disease-oriented research. Within this dataset, 223 rare exome mutations in the ZPR1 gene (ZEMs) were identified across these six groups, showing a minor allele frequency (MAF) ranging from 3.98e-08 to 0.0658. Notably, only two of these mutations, A419V (rs144966144) and I196T (rs368697578), are listed in the ClinVar database [32]. These rare ZEMs were found in 344 individuals as heterozygous carriers and 5 as homozygous.

We analyzed the 223 rare ZEMs by calculating their SIFT scores and PolyPhen-2_HVAR scores. According to our findings (see Supplementary Information 1), 60 of these mutations were identified as pathogenic. Notably, the I196T mutation (rs368697578) that was first found in ClinVar was correctly identified as harmful by both SIFT and PolyPhen-2_HVAR. The A419V mutation (rs144966144), which was also found in ClinVar annotated as benign and in conflict with these results. The results also indicated that the PolyPhen-2_HVAR score of 0.729, which means it might be harmful, but a SIFT score of 0.46 meant it was probably tolerated. As a result, PolyPhen-2_HVAR correctly identified this mutation while SIFT did not. Beyond these, both SIFT and PolyPhen-2 categorized 119 mutations as tolerated. The remaining 44 rare ZEMs could not be conclusively classified and therefore excluded from further analysis.

To discern between pathogenic and tolerated mutations in the ZPR1 gene, we employed ten additional algorithms, with the findings detailed in Supplementary Information (SI) sections 3 and 4. Figure 1 in our report illustrates the location of each amino acid in the ZPR1 protein, including the N-disordered region (amino acids 1–32), ZnF1 (51–83), and ZnF2 (259–291) regions. The protein is further divided into two domains: the A-domain (amino acids 100–234) and the B-domain (amino acids 308–438), as previously outlined [4]. Notably, a number of pathogenic ZPR1 exome mutations (ZEMs) (25 out of 60) are located within the A-domain, which is known for its interaction with eEF1A. The B-domain contains the second highest number of pathogenic ZEMs (15 out of 60) (SI 8). The mutation I196T (rs368697578), associated with a rare ZPR1 disorder [4], is situated in the A-domain. In terms of pathogenic ZEMs distribution, the highest count is in the region encoding the A-domain (236 out of 349 individuals), followed by mutations in the B-domain (26 individuals), the ZnF1 and ZnF2 domains (16 individuals), and other areas (16 individuals), as elaborated in SI 8.

**Fig 1.**
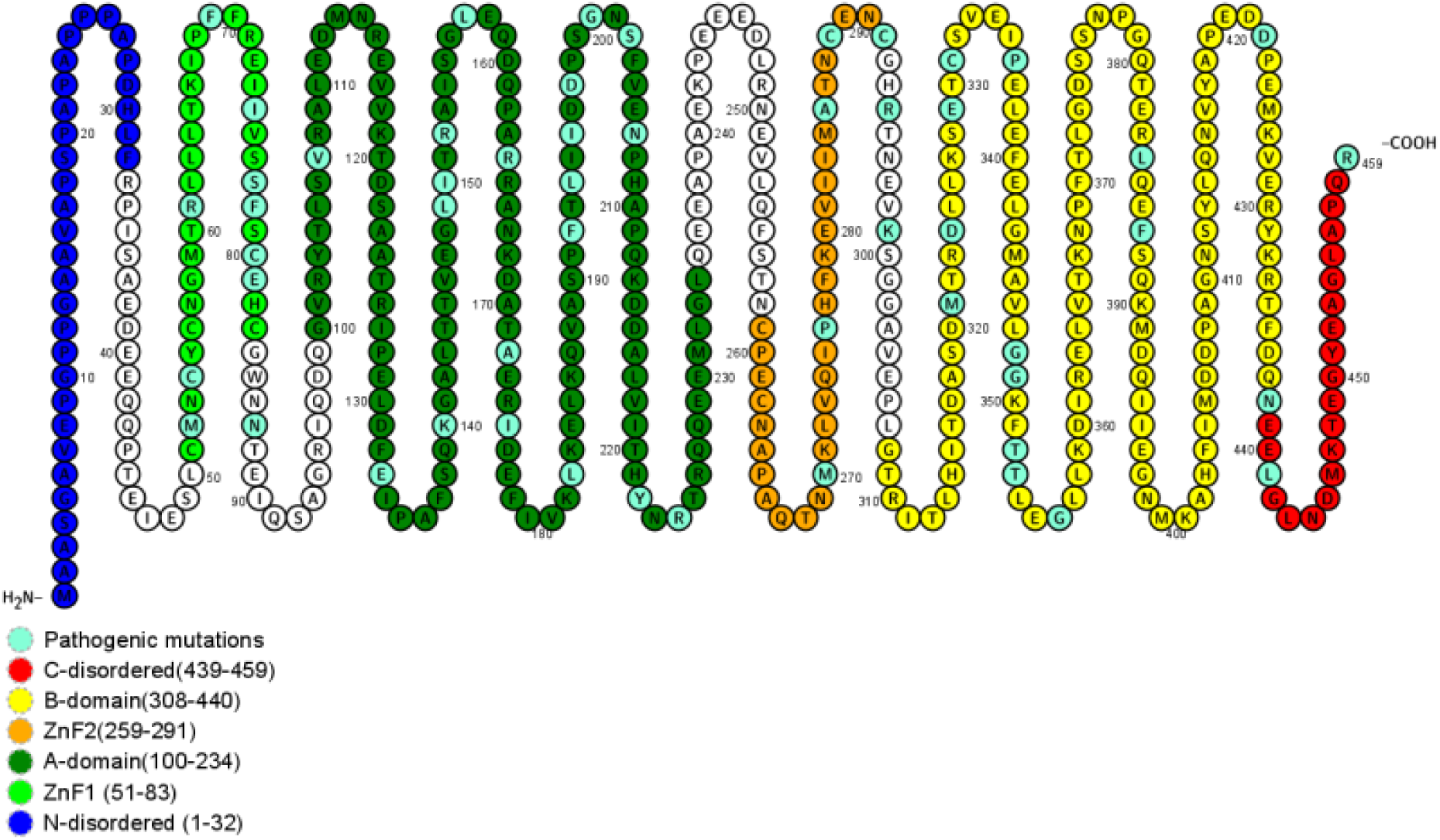
ZPR1 protein model generated with the software PROTTER [57]. This model indicates the locations of ancestral amino acids, including the location of the pathogenic ZEMs.

In summary, in this analysis, the gnomAD data containing the information from 122,678 individuals was sampled. 223 ZEMs were found in 349 individuals. 60 of these rare ZEMs were classified as pathogenic, and 119 were classified as tolerated. In another prediction, the effects of the classified mutations were made clearer. Looking at all those who were sampled, 90% of the pathogenic ZEMs (54/60) are found in 44/122,678 of the population, while 10% (6/60) are found in 227/122,678.

### Ethnic differences in the prevalence of ZPR1 pathogenic mutations

The prevalence of heterozygous and homozygous individuals carrying the identified ZPR1 pathogenic mutations are shown in Table 1 by ethnic groups. Also, the data for the neutral mutations is shown in SI (4).

**Table 1.** Ethnic differences and global prevalence of pathogenic ZPR1 mutations.

|  | Global<br>(N = 122,678) | African/<br>African<br>American<br>(N = 8,128) | Latino/<br>American<br>(N = 17,296) | Ashkenazi<br>Jews<br>(N = 5,040) | East Asian<br>(N = 9,197) | European<br>(N = 67,709) | South Asian<br>(N = 15,308) |
| --- | --- | --- | --- | --- | --- | --- | --- |
| <sup>a</sup> Total<br>Alleles, N | 245,356 | 16,256 | 34,592 | 10,080 | 18,394 | 135,418 | 30,616 |
| Heterozygous<br>Carriers, N | 330 | 28 | 50 | 2 | 6 | 217 | 27 |
| % | 0.27 | 0.34 | 0.29 | 0.04 | 0.07 | 0.32 | 0.18 |
| Homozygous<br>Carriers, N | 3 | 0 | 0 | 0 | 0 | 1 | 2 |
| % | 0.0024 | 0 | 0 | 0 | 0 | 0.0015 | 0.013 |
| <sup>b</sup> Mutated<br>Alleles, N | 336 | 28 | 50 | 2 | 6 | 219 | 31 |
| % | 0.0014 | 0.34 | 0.39 | 0.04 | 0.07 | 0.32 | 0.20 |
| **Ethnic-**<br>**Specific**<br>**Mutations** |  |  |  |  |  |  |  |
| Heterozygous<br>Carriers, N | 65 | 10 | 3 | 1 | 0 | 33 | 18 |
| Homozygous<br>Carriers, N | 2 | 0 | 0 | 0 | 0 | 0 | 2 |
| Mutated<br>Alleles | 69 | 10 | 3 | 1 | 0 | 33 | 22 |

<sup>a</sup> The total number of alleles in each ethnic group, determined by multiplying the number of individuals by two.
<sup>b</sup> The pathogenic alleles in each ethnic group.

**Table 2.** Ethnic differences and global prevalence of neutral ZPR1 mutations.

|  | Global<br>(N =122,678) | African/<br>American<br>(N =8,128) | Latino/<br>American<br>(N =17,296) | Ashkenazi<br>Jews<br>(N = 5,040) | East Asian<br>(N = 9,197) | European<br>(N= 67,709) | South Asian<br>(N = 15,308) |
| --- | --- | --- | --- | --- | --- | --- | --- |
| Total Alleles,<br>N | 245,356 | 16,256 | 34,592 | 10,080 | 18,394 | 135,418 | 30,616 |
| Heterozygous<br>Carriers, N | 21,476 | 573 | 6,892 | 971 | 105 | 11,528 | 1,407 |
| % | 17.51 | 7.05 | 39.85 | 19.27 | 1.14 | 17.03 | 9.19 |
| Homozygous<br>Carriers, N | 806 | 11 | 438 | 30 | 0 | 301 | 27 |
| % | 0.66 | 0.14 | 2.53 | 0.59 | 0 | 0.44 | 0.18 |
| Mutated<br>Alleles, N | 23,088 | 595 | 7,768 | 1,031 | 105 | 12,130 | 1,461 |
| % | 9.41 | 3.66 | 22.46 | 10.23 | 0.57 | 8.96 | 4.77 |
| **Ethnic-**<br>**Specific**<br>**Mutations** |  |  |  |  |  |  |  |
| Heterozygous<br>Carriers, N | 176 | 17 | 33 | 1 | 23 | 73 | 29 |
| Homozygous<br>Carriers, N | 36 | 0 | 35 | 1 | 0 | 0 | 0 |
| Mutated<br>Alleles | 248 | 17 | 103 | 3 | 23 | 73 | 29 |

Eight pathogenic mutations were found in the gnomAD dataset, which has 122,678 people. These include p.Asp438Ile and p.Thr353Ile in the A-domain and p.Asp207Lys in the B-domain. These mutations are present in only three homozygous individuals (0.00245%, 3/122,678), as depicted in Figure 2, which breaks down the data by ethnic groups. Notably, of the mutations observed in at least five individuals, p.Asp438Ile was exclusively found in males. According to Table 1, 330 heterozygous carriers (0.27%, 330/122,678) possess the remaining 52 pathogenic mutations. This includes 217 Europeans (0.32%, 217/67,709), 50 Latino Americans (0.29%, 50/17,296), 27 South Asians (0.18%, 27/15,308), six East Asians (0.07%, 6/9197), 28 African/African Americans (0.34%, 28/8128), and two Ashkenazi Jews (0.04%, 2/5040). There are big differences between the six racial and ethnic groups in how these harmful ZEMs are spread. For example, the Ashkenazi Jewish group has the lowest rate (0.04%), while the African American group has the highest rate (0.34%).

**Fig 2.**
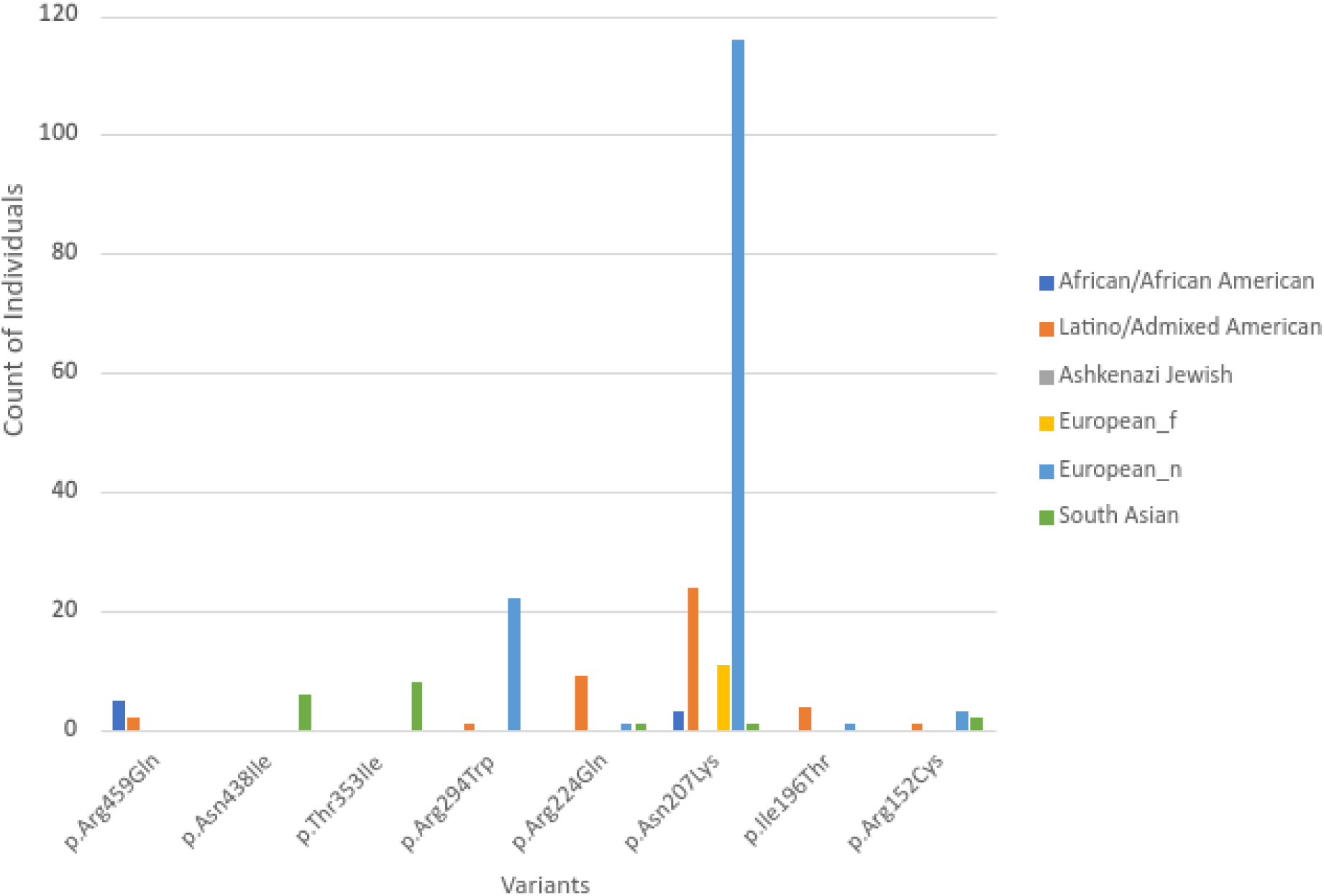
The distribution of pathogenic variants among different ethnic groups in at least five individuals.

In summary, out of the 60 pathogenic mutations identified, 28 were exclusive to Europeans, three to Latino/Americans, eight to South Asians, and seven to African/African Americans, while none were specific to Ashkenazi Jewish or East Asian populations (as shown in Table 1). Additionally, 46 of these pathogenic mutations displayed an ethnic-specific pattern and found in 67 heterozygous carriers and 2 homozygous carriers. Notably, both of the homozygous carriers were of South Asian descent.

### Evolutionary conservation analysis of the ZPR1 protein

Mutations that destabilize proteins often result in degradation by the proteasome, leading to reduced protein levels in cells [45, 46], and can also cause misfolding and/or aggregation [48]. Furthermore, disturbances in protein stability may act as a contributing factor in diseases linked to genes with haploinsufficiency [47]. The relationship between the severity of diseases and structurally destabilizing mutations in genetic or monogenic disorders has been substantiated by Scheller et al. [46] and Yue et al. [45]. To investigate to what extent the 179, including 60 pathogenic and 119 tolerated, rare ZEMs, affect stability of the protein structure, we calculated the difference in free energy of folding (ΔΔG in kcal/mol) between a wild-type ZPR1 residue and the corresponding mutant, i.e., ΔΔG = ΔGmut - ΔGwt, using FoldX [49], an algorithm that evaluates the effects of mutations using an empirical force field where the side chains are allowed to move but not the backbone. A positive ΔΔG indicates the destabilization effect, while a negative indicates stabilization. The experimental error twilight zone is ΔΔG = ± 0.5 kcal/mol, so mutations with the calculated ΔΔG = ± 0.5 kcal/mol are defined as twilight mutations [50]. The same metric was adopted in our test. Also, mutations with ΔΔG up to 1 kcal/mol have been described as “mild destabilizers” [50]. This metric was adopted too. Mutants with ΔΔG > 2 kcal/mol were found to make the folded state less stable, which could not be told apart from mutants that cause disease [51]. In our test, a mutation is considered significant if ΔΔG > 1.5 kcal/mol. Residues with ΔΔG > 1.5 kcal/mol were identified and shown in Figure 3.

**Fig 3.**
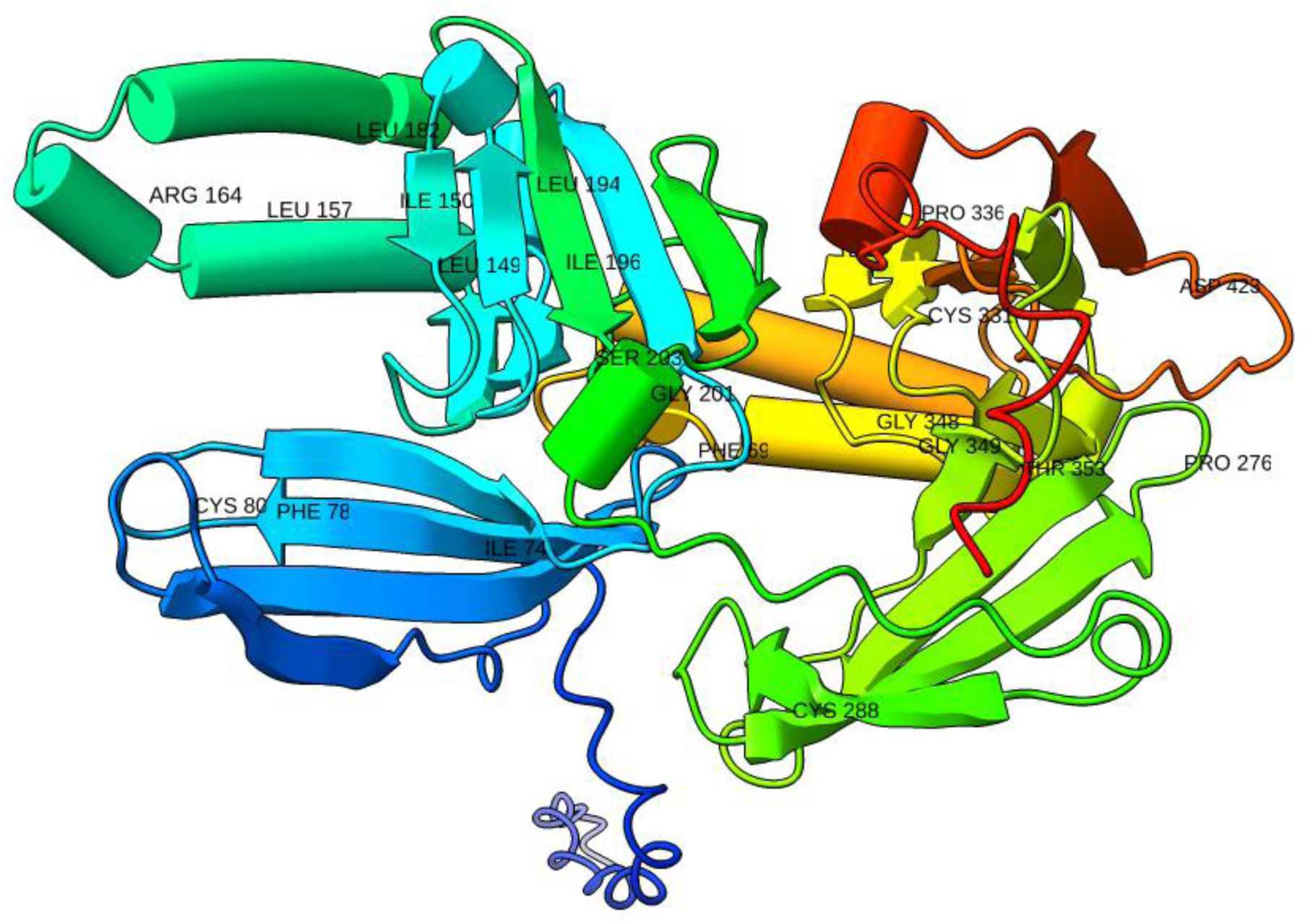
The 3D structure model for ZPR1 protein (Uniprot code: AF-O75312-F1). The structure is colored from N-ter (blue) to C-ter (red). The mutated residues with ΔΔG >= 1.5 kcal/mol are labeled. The figure was generated using ChimeraX [58]. The color range is determined by five color wells, which correspond to the starting color, three intermediate interpolation colors, and the ending color from blue (N-terminal), cyan, green, yellow, and red (C-terminal), in that order. **α**-helices = cylinders **β**-strands = arrows where arrowheads points in the N-terminus to C-terminus direction non alpha/beta structures = threads or strings

The calculated ΔΔG values ranged from −0.95 kcal/mol (p.Ser77Tyr) to −0.003 kcal/mol (p.Tyr222Cys) for the pathogenic mutations and from −1.453 kcal/mol (p.His293Tyr) to −0.008 kcal/mol for the neutral mutations. Specifically, 23 mutations with ΔΔG > 1.5 kcal/mol were thus considered to cause significant structural changes. Of these 23, fourteen affected the structural residues G349, C331, C288, I196, L194, L182, L157, I150, L149S, C80, F78, I74, and F69, and nine affected the functional residues D423, T353, G348, P336, P276, S203, G201, and R164. Based on recommendations from the previous studies [52], a residue at a position that is highly conserved and exposed was proposed to be functional; a residue at a position that is highly conserved but buried was proposed to be structural. In the ZPR1 protein, residue at position 196 is buried in the sixth beta-strand of the A-domain that occupies the hydrophobic core [4]. The hydrophobic core network consists of residues I196, L149, L194, A123, V117, Y103, and V205 [4]. A mutation changing this residue will disrupt this network [4]. In our test, a ΔΔG of 2.3 kcal/mol, indicating a disease-causing mutant, was obtained for mutation p.IIe196Thr. Using ConSurf [28], we confirmed that the residue at the position of 196 is buried and structurally important (SI (7)). A previous study demonstrated that the mutation p.Ile196Thr impedes cell cycle progression beyond the G1 phase [4]. In comparison to control fibroblast cells, hardly any cells from affected patients managed to progress through the cell cycle, and only a minor fraction of the patient’s cells was observed in the late S phase and G2/M phases [4]. This residue is found in the ZnF1-A domain of the ZPR1 protein. This domain is particularly important for binding eEF1A, which is needed for normal cell growth, proliferation, and cycle progression. Therefore, the mutation p.Ile196Thr might play a significant role in the interaction with eEF1A [14].

We used ConSurf to find changes in buried ZnF1-A domain residues that are very likely to disrupt the network of interactions between residues. They are (with the ΔΔG values in kcal/mol shown in parentheses): p.Leu194Pro (2.4), p.Leu194Gln (2.2), p.Leu182Pro (5.3), p.Leu157Pro (2.3), p.Ile150Thr (1.9), p.Leu149S (2.6), p.Cys80Arg (4.1), p.Phe78Cys (3.2), p.Ile74Thr (2.0), and p.Phe69Cys (2.3). Besides, the substitutions p.Ser203Gly (1.8), p.Gly201Arg (9.6), p.Arg164Pro (2.8), and p.Arg164Gly (1.6) that affect exposed residues were also identified. The substitutions p.Gly349Asp (3.6), p.Cys331Ser (1.8), p.Cys288Ser (1.6), p.Asp423Val (2.3), p.Thr353Ile (1.76), p.Gly348Arg (13.1), p.Pro336Leu (2.0), and p.Pro276Ser (1.6) could affect both structural and exposed residues, which could cause changes in the structure. These mutations, whether buried or exposed, were determined to significantly impact the structure. It was discovered that the following mutations could change structural residues and exposed residues that could change structure: p.Gly349Asp (3.6), p.Cys331Ser (1.8), p.Cys288Ser (1.6), p.Asp423Val (2.3), p.Thr353Ile (1.76), p.Gly348Arg (13.1), p.Pro336Leu (2.0), and p.Pro276Ser (1.6), whether they were buried or not.

In the case of a monomeric protein, FoldX predicts its overall stability by considering a range of energy contributions. These include backbone and side chain hydrogen bonds, van der Waals and electrostatic interactions, penalties for burying polar and hydrophobic groups, inter-residue van der Waals clashes, entropic costs for fixing side chains and the main chain, the energy cost of cis peptide bonds, intra-residue van der Waals torsional clashes, electrostatics from helix dipoles, water bridges, disulfide bonds, partial covalent bonds (such as metal interactions), ionization energy, and the count of residues [49]. The side chain atoms of the mutated residue play a notable role in ΔΔG calculation [49]. Utilizing Python 3.10.2 [53], we analyzed the Pearson correlation between ΔΔG and various FoldX energy terms. We discovered a moderate correlation with torsional clashes (from intra-residue van der Waals torsional clashes; Pearson correlation 0.49); and a strong correlation with van der Waals clashes (Pearson correlation 0.86). In terms of solvation polarity, the correlation was good (0.41). Our data indicated that torsional and van der Waals clashes had more influence on ΔΔG than other factors. This observation, where one or two energy terms provide a clearer trend than the total energy, aligns with findings from a previous study on laccase mutants [54]. Overall, our FoldX results are consistent with prior experimental findings for ZPR1, except for those related to the disordered region [49]. This accounts for the high ΔΔG value (20.2 kcal/mol) for the A21V mutation, which occurs in the disordered region.

## Discussion

This study analyzed 223 rare ZEMs, all with low minor allele frequencies (MAF) ranging from 3.98e-8 to 0.0658. A major portion (94.1%) had MAFs between 3.98e-06 and 9.55e-05. Despite previous suggestions [55] that pathogenicity is not always tied to extremely low frequencies, the mutation p.Ala264Val (rs35120633) associated with plasma triglyceride levels [36] and identified as pathogenic in our study had the highest MAF at 0.0658. Ethnic disparities were evident among the 60 pathogenic ZEMs identified, with over 80% found in the African/African American population (SI 3). Three homozygous carriers of rare ZEMs were detected, two in South Asians and one in a European, possibly indicating late-onset or non-fully penetrant diseases [37]. Interestingly, the ZPR1 gene was not mentioned in ORPHANET [56], the portal for rare diseases and orphan drugs, as of January 10, 2022.

Our analysis examined the prevalence of ZPR1 mutations across various ethnic groups using data from gnomAD, the largest exome and genome sequencing database available. This approach minimized bias [38] and provided significant statistical results [39], though it lacked representation from Arab, Pacific Island, and Native Australian groups [37]. The study underlines the need for further clinical and in vitro functional characterizations to confirm the pathogenic nature of these mutations.

We assessed the impact of amino acid substitutions from the 223 rare ZEMs on protein stability, showing their positions in the ZPR1 protein domains (Fig. 1) and providing a basis for future functional analysis. Using FoldX software, we calculated the ΔΔG for these mutations. Although FoldX is highly regarded for identifying pathogenic mutations and efficient in performance [51], caution is advised in interpreting its results, especially in disordered regions. Our results showed that more than half of the ACMG/ACP variants changed the structure of the protein. This shows that predicted structural changes and VEP algorithms are similar.

This study meets the ACMG/AMP evidence standards by putting all exome variants in ZPR1 from the gnomAD database into two groups: those that cause disease and those that don’t. We used SIFT and PolyPhen-2-HVAR consensus to find 60 pathogenic, 119 neutral, and 44 variants of unknown significance (VUS). The increased use of multiple VEPs, however, showed limitations in reducing VUS, particularly for variants not previously associated with disease [53]. The frequency of ZPR1 exome variants challenges the traditional understanding of genetic causality based on MAF. We saw a strong link between ΔΔG and steric clashes. It is important to note that variants with a ΔΔG of at least ±1.5 kcal/mol require more research (SI 6). The study highlights the importance of genetic analysis before marriage as a preventive measure against monogenic disorders and ARD-related burdens. Our data can be a valuable resource for population genetic screening programs, especially in studying autosomal recessive disorders that are typically under-researched.

## Acknowledgements

This research was supported by the National Institutes of Health (NIH) Common Fund (U24 CA224370), UNM Comprehensive Cancer Center Support Grant NCI (P30CA118100), and an Institutional Development Award (IDeA) from the National Institute of General Medical Sciences of the National Institutes of Health (P20GM103451).

## Supporting Information

All the relevant data, the codes, and the corresponding supporting files are available at https://github.com/abok2020/Evaluation-of-ZPR1-exome-variants/blob/main/supplementary_data.xlsx https://github.com/abok2020on GitHub.

## References

1. UniProt Consortium. UniProt: the universal protein knowledgebase in 2021. Nucleic Acids Res [Internet].2021;49(D1):D480–9. Available from: 10.1093/nar/gkaa1100

2. Chittilla M, Akimbekov NS, Razzaque MS. High-fat diet-associated cognitive decline: Is zinc finger protein 1 (ZPR1) the molecular connection? Curr Res Physiol. 2021;4:223–8. 10.1016/j.crphys.2021.09.004

3. Hamosh A, Scott AF, Amberger JS, Bocchini CA, McKusick VA. Online Mendelian Inheritance in Man (OMIM), a knowledgebase of human genes and genetic disorders. Nucleic Acids Res. 2005;33(Database issue):D514–7. 10.1093/nar/gki033

4. Ito YA, Smith AC, Kernohan KD, Pena IA, Ahmed A, McDonell LM, et al. A ZPR1 mutation is associated with a novel syndrome of growth restriction, distinct craniofacial features, alopecia, and hypoplastic kidneys. Clin Genet. 2018;94(3–4):303–12. 10.1111/cge.13388

5. Wall JD, Sathirapongsasuti JF, Gupta R, Rasheed A, Venkatesan R, Belsare S, et al. South Asian medical cohorts reveal strong founder effects and high rates of homozygosity. Nat Commun. 2023;14(1):3377. 10.1038/s41467-023-38766-1

6. Lanzinger M. Consanguineous marriages: Contexts and controversies. In: Administrating Kinship: Marriage Impediments and Dispensation Policies in the 18th and 19th Centuries. Brill | Nijhoff; 2023. p. 280–344.

7. Klingseisen A, Jackson AP. Mechanisms and pathways of growth failure in primordial dwarfism. Genes Dev. 2011;25(19):2011–24. 10.1101/gad.169037

8. Bicknell LS, Walker S, Klingseisen A, Stiff T, Leitch A, Kerzendorfer C, et al. Mutations in ORC1, encoding the largest subunit of the origin recognition complex, cause microcephalic primordial dwarfism resembling Meier-Gorlin syndrome. Nat Genet. 2011;43(4):350–5. 10.1038/ng.776

9. Bicknell LS, Bongers EMHF, Leitch A, Brown S, Schoots J, Harley ME, et al. Mutations in the pre-replication complex cause Meier-Gorlin syndrome. Nat Genet. 2011;43(4):356–9. 10.1038/ng.775

10. Mokrani-Benhelli H, Gaillard L, Biasutto P, Le Guen T, Touzot F, Vasquez N, et al. Primary microcephaly, impaired DNA replication, and genomic instability caused by compound heterozygous ATR mutations. Hum Mutat. 2013;34(2):374–84. 10.1002/humu.22245

11. O’Driscoll M, Ruiz-Perez VL, Woods CG, Jeggo PA, Goodship JA. A splicing mutation affecting expression of ataxia-telangiectasia and Rad3-related protein (ATR) results in Seckel syndrome. Nat Genet. 2003;33(4):497–501. 10.1038/ng1129

12. Gangwani L, Mikrut M, Galcheva-Gargova Z, Davis RJ. Interaction of ZPR1 with translation elongation factor-1α in proliferating cells. J Cell Biol. 1998;143(6):1471–84. 10.1083/jcb.143.6.1471

13. Galcheva-Gargova Z, Gangwani L, Konstantinov KN, Mikrut M, Theroux SJ, Enoch T, et al. The cytoplasmic zinc finger protein ZPR1 accumulates in the nucleolus of proliferating cells. Mol Biol Cell. 1998;9(10):2963–71. 10.1091/mbc.9.10.2963

14. Galcheva-Gargova Z, Konstantinov KN, Wu IH, Klier FG, Barrett T, Davis RJ. Binding of zinc finger protein ZPR1 to the epidermal growth factor receptor. Science. 1996;272(5269):1797–802. 10.1126/science.272.5269.1797

15. Doran B, Gherbesi N, Hendricks G, Flavell RA, Davis RJ, Gangwani L. Deficiency of the zinc finger protein ZPR1 causes neurodegeneration. Proc Natl Acad Sci USA. 2006;103(19):7471–5. 10.1073/pnas.0602057103

16. Gangwani L. Deficiency of the zinc finger protein ZPR1 causes defects in transcription and cell cycle progression. J Biol Chem. 2006;281(52):40330–40. 10.1074/jbc.M608165200

17. Mishra AK, Gangwani L, Davis RJ, Lambright DG. Structural insights into the interaction of the evolutionarily conserved ZPR1 domain tandem with eukaryotic EF1A, receptors, and SMN complexes. Proc Natl Acad Sci USA. 2007;104(35):13930–5. 10.1073/pnas.0704915104

18. Yanaka N, Kaseda Y, Tanaka A, Nogusa Y, Sumiyoshi N, Kato N. Generation of a zinc finger protein ZPR1 mutant that constitutively interacted with translation elongation factor 1alpha. Biosci Biotechnol Biochem. 2009;73(12):2809–11. 10.1271/bbb.90745

19. Abbott CM, Newbery HJ, Squires CE, Brownstein D, Griffiths LA, Soares DC. eEF1A2 and neuronal degeneration. Biochem Soc Trans. 2009;37(Pt 6):1293–7. 10.1042/BST0371293

20. Kannan A, Jiang X, He L, Ahmad S, Gangwani L. ZPR1 prevents R-loop accumulation, upregulates SMN2 expression and rescues spinal muscular atrophy. Brain. 2020;143(1):69–93. 10.1093/brain/awz373

21. Guan F, Niu Y, Zhang T, Liu S, Ma L, Qi T, et al. Two-stage association study to identify the genetic susceptibility of a novel common variant of rs2075290 in ZPR1 to type 2 diabetes. Sci Rep. 2016;6(1). 10.1038/srep29586

22. Bai W, Kou C, Zhang L, You Y, Yu W, Hua W, et al. Functional polymorphisms of the APOA1/C3/A4/A5-ZPR1-BUD13 gene cluster are associated with dyslipidemia in a sex-specific pattern. PeerJ. 2019;6(e6175):e6175. 10.7717/peerj.6175

23. Liu B, Xing X, Li X, Guo Q, Xu T, Xu K. ZNF259 promotes breast cancer cells invasion and migration via ERK/GSK3β/snail signaling. Cancer Manag Res. 2018;10:3159–68. 10.2147/CMAR.S174745

24. Shan Y, Cao W, Wang T, Jiang G, Zhang Y, Yang X. ZNF259 inhibits non-small cell lung cancer cells proliferation and invasion by FAK-AKT signaling. Cancer Manag Res. 2017;9:879–89. 10.2147/cmar.s150614

25. Wang K, Li M, Hakonarson H. ANNOVAR: functional annotation of genetic variants from high-throughput sequencing data. Nucleic Acids Res. 2010;38(16):e164. 10.1093/nar/gkq603

26. Ng PC, Henikoff S. SIFT: Predicting amino acid changes that affect protein function. Nucleic Acids Res. 2003;31(13):3812–4. 10.1093/nar/gkg509

27. Adzhubei IA, Schmidt S, Peshkin L, Ramensky VE, Gerasimova A, Bork P, et al. A method and server for predicting damaging missense mutations. Nat Methods. 2010;7(4):248–9. 10.1038/nmeth0410-248

28. Ashkenazy H, Abadi S, Martz E, Chay O, Mayrose I, Pupko T, et al. ConSurf 2016: an improved methodology to estimate and visualize evolutionary conservation in macromolecules. Nucleic Acids Res. 2016;44(W1):W344–50. 10.1093/nar/gkw408

29. Berezin C, Glaser F, Rosenberg J, Paz I, Pupko T, Fariselli P, et al. ConSeq: the identification of functionally and structurally important residues in protein sequences. Bioinformatics. 2004;20(8):1322–4. 10.1093/bioinformatics/bth070

30. Yang J, Zhang Y. Protein structure and function prediction using I-TASSER. Curr Protoc Bioinformatics. 2015;52(1):5.8.1–5.8.15. 10.1002/0471250953.bi0508s52

31. Berman HM. The Protein Data Bank. Nucleic Acids Res. 2000;28(1):235–42. 10.1093/nar/28.1.235

32. Landrum MJ, Lee JM, Benson M, Brown GR, Chao C, Chitipiralla S, et al. ClinVar: improving access to variant interpretations and supporting evidence. Nucleic Acids Res. 2018;46(D1):D1062–7. 10.1093/nar/gkx1153

33. Jagadeesh KA, Wenger AM, Berger MJ, Guturu H, Stenson PD, Cooper DN, et al. M-CAP eliminates a majority of variants of uncertain significance in clinical exomes at high sensitivity. Nat Genet. 2016;48(12):1581–6. 10.1038/ng.3703

34. Feng B-J. PERCH: A unified framework for disease gene prioritization. Hum Mutat. 2017;38(3):243–51. 10.1002/humu.23158

35. Alirezaie N, Kernohan KD, Hartley T, Majewski J, Hocking TD. ClinPred: Prediction tool to identify disease-relevant nonsynonymous single-nucleotide variants. Am J Hum Genet. 2018;103(4):474–83. 10.1016/j.ajhg.2018.08.005

36. Shihab HA, Rogers MF, Gough J, Mort M, Cooper DN, Day INM, et al. An integrative approach to predicting the functional effects of non-coding and coding sequence variation. Bioinformatics. 2015;31(10):1536–43. 10.1093/bioinformatics/btv009

37. Chiorean A, Garver WS, Meyre D. Signatures of natural selection and ethnic-specific prevalence of NPC1 pathogenic mutations contributing to obesity and Niemann-Pick disease type C1. Sci Rep. 2020;10(1):18787. Available from: 10.1038/s41598-020-75919-4

38. Karczewski KJ, Francioli LC, Tiao G, Cummings BB, Alföldi J, Wang Q, et al. The mutational constraint spectrum quantified from variation in 141,456 humans. Nature. 2020;581(7809):434–43. 10.1038/s41586-020-2308-7

39. Pirastu N, Cordioli M, Nandakumar P, Mignogna G, Abdellaoui A, Hollis B, et al. Genetic analyses identify widespread sex-differential participation bias. Nat Genet. 2021;53(5):663–71. 10.1038/s41588-021-00846-7

40. Vaser R, Adusumalli S, Leng SN, Sikic M, Ng PC. SIFT missense predictions for genomes. Nat Protoc. 2016;11(1):1–9. 10.1038/nprot.2015.123

41. Chun S, Fay JC. Identification of deleterious mutations within three human genomes. Genome Res [Internet]. 2009;19(9):1553–61. 10.1101/gr.092619.109

42. Schwarz JM, Cooper DN, Schuelke M, Seelow D. MutationTaster2: mutation prediction for the deep-sequencing age. Nat Methods. 2014;11(4):361–2. 10.1038/nmeth.2890

43. Reva B, Antipin Y, Sander C. Predicting the functional impact of protein mutations: application to cancer genomics. Nucleic Acids Res. 2011;39(17):e118. 10.1093/nar/gkr407

44. Choi Y, Chan AP. PROVEAN web server: a tool to predict the functional effect of amino acid substitutions and indels. Bioinformatics. 2015;31(16):2745–7. 10.1093/bioinformatics/btv195

45. Yue P, Li Z, Moult J. Loss of protein structure stability as a major causative factor in monogenic disease. J Mol Biol. 2005;353(2):459–73. 10.1016/j.jmb.2005.08.020

46. Scheller R, Stein A, Nielsen SV, Marin FI, Gerdes A-M, Di Marco M, et al. Toward mechanistic models for genotype-phenotype correlations in phenylketonuria using protein stability calculations. Hum Mutat. 2019;40(4):444–57. 10.1002/humu.23707

47. Birolo G, Benevenuta S, Fariselli P, Capriotti E, Giorgio E, Sanavia T. Protein stability perturbation contributes to the loss of function in haploinsufficient genes. Front Mol Biosci. 2021;8:620793. 10.3389/fmolb.2021.620793

48. Ajmal MR. Protein misfolding and aggregation in proteinopathies: Causes, mechanism and cellular response. Diseases. 2023;11(1). 10.3390/diseases11010030

49. Schymkowitz J, Borg J, Stricher F, Nys R, Rousseau F, Serrano L. The FoldX web server: an online force field. Nucleic Acids Res. 2005;33(Web Server issue):W382–8. 10.1093/nar/gki387

50. Marabotti A, Del Prete E, Scafuri B, Facchiano A. Performance of Web tools for predicting changes in protein stability caused by mutations. BMC Bioinformatics. 2021;22(Suppl 7):345. 10.1186/s12859-021-04238-w

51. Randles LG, Lappalainen I, Fowler SB, Moore B, Hamill SJ, Clarke J. Using model proteins to quantify the effects of pathogenic mutations in Ig-like proteins. J Biol Chem. 2006;281(34):24216–26. 10.1074/jbc.M603593200

52. Guharoy M, Chakrabarti P. Conserved residue clusters at protein-protein interfaces and their use in binding site identification. BMC Bioinformatics. 2010;11(1):286. 10.1186/1471-2105-11-286

53. Van Rossum G, Drake FL. Python 3 Reference Manual. Scotts Valley, CA: CreateSpace; 2009.

54. Christensen NJ, Kepp KP. Accurate stabilities of laccase mutants predicted with a modified FoldX protocol. J Chem Inf Model. 2012;52(11):3028–42. 10.1021/ci300398z

55. Lek M, Exome Aggregation Consortium, Karczewski KJ, Minikel EV, Samocha KE, Banks E, et al. Analysis of protein-coding genetic variation in 60,706 humans. Nature. 2016;536(7616):285–91. 10.1038/nature19057

56. Slebodnik M. Orphanet: The Portal for Rare Diseases and Orphan Drugs2009384Orphanet: The Portal for Rare Diseases and Orphan Drugs. Paris: Institute National de la Santé et de la Recherche Médicale (INSERM) Last visited June 2009. Gratis URL: www.orpha.net/. Ref Rev. 2009;23(8):45–6. 10.1108/09504120911003492

57. Omasits U, Ahrens CH, Müller S, Wollscheid B. Protter: interactive protein feature visualization and integration with experimental proteomic data. Bioinformatics. 2014;30(6):884–6. 10.1093/bioinformatics/btt607

58. Pettersen EF, Goddard TD, Huang CC, Meng EC, Couch GS, Croll TI, et al. UCSF ChimeraX: Structure visualization for researchers, educators, and developers. Protein Sci. 2021;30(1):70–82. 10.1002/pro.3943

